# Did variants in inborn errors of immunity genes contribute to the extinction of Neanderthals?

**DOI:** 10.1101/2022.10.19.512108

**Authors:** Zijun Zhou, Sigrid M.A Swagemakers, Mirthe S. Lourens, Narissara Suratannon, Peter J. van der Spek, Virgil A.S.H. Dalm, Willem A. Dik, Hanna IJspeert, P. Martin van Hagen

## Abstract

Neanderthals were a species of archaic human that became extinct around 40,000 years ago. Modern humans have inherited 1-6% of Neanderthal DNA as a result of interbreeding with the Neanderthals. These inherited Neanderthal genes have paradoxical influences, while some can provide protection to viral infections, some others are associated with autoimmune/auto-inflammatory diseases.

We hypothesized that genetic variants with strong detrimental effects on the function of the immune system could potentially contributed to the extinction of the Neanderthal population. In modern humans more than 450 genes are associated with inborn errors of immunity (IEI). We used the publically available genome information from a Neanderthal from the Altai mountains and filtered for potentially damaging variants that were present in genes associated with IEI, and checked whether these variants were present in the genomes of the Denisovan, Vindija and Chagyrskaya Neanderthals.

We identified 24 homozygous variants and 15 heterozygous variants in IEI-related genes in the Altai Neanderthal. Interestingly, two homozygous variants in the *UNC13D* gene and one variant in the *MOGS* gene were present in all archaic genomes. Defects in the *UNC13D* gene are known to cause a severe and often fatal disease called hemophagocytic lymphohistiocystosis (HLH). One of these variants p.(Asn943Ser) has been reported in patients with HLH. Variants in *MOGS* are associated with glycosylation defects in the immune system affecting the susceptibility for infections. So, although we do not know exactly the functional impact yet, these three variants could have resulted in an increased susceptibility to severe diseases, and may have contributed to the extinction of Neanderthals after exposure to specific infections.

## Introduction

Neanderthals were a group of archaic humans that used to inhabit Eurasia. Neanderthals lived in relatively small social groups (10-30 individuals) (*1*), probably composed of individuals of both sexes and different age classes (*1*). A related but distinctive sister group of Neanderthals known as the Denisovans lived at the same time in Siberia and eastern Asia (*2*). Ancestors of the Neanderthals and Denisovans diverged from the main human lineage at least 600,000 years ago, and then split from each other around 400,000 years ago (*3–6*). Although they lived mainly in isolated groups, they were not completely isolated from the early modern humans that migrated from Africa (*3, 7*). Archaeological evidence support that these groups of archaic humans and early modern humans shared the same habitat until, by some yet-unknown cause, Neanderthals and Denisovans disappeared around 40,000 years ago. Their extinction coincided with the migration of early modern humans outside of Africa starting around 50,000 years ago (*8*). Early modern humans have co-existed with Neanderthals and Denisovans for at least 10,000 years which allowed for interbreeding, even though of low magnitude between these two groups (*9*). Gene flow analysis showed that genomes of modern Eurasians contain around 1 – 6% DNA inherited from interbreeding with Neanderthals and Denisovans (*3, 9–12*).

Although the exact function of the surviving archaic human genes in modern humans genomes remains unclear, several recent genomic studies suggested that the inherited archaic alleles may explain certain human clinical traits (*13, 14*). Genetic variants impacting immunity have been shown to be transferred from archaic humans to modern humans (*15–17*). In particular, innate immunity regulatory genes as the *TLR6-1-10* gene cluster and the *MERTK* gene were introduced into modern human European genomes from both Neanderthals and Denisovans (*16*). Furthermore, it is suggested that alleles encoding for proteins which provide protection against RNA viruses in modern Europeans are most likely inherited from the Neanderthals (*18, 19*). The conservation of such alleles indicates that they could have been protective to both modern humans and Neanderthals when they were exposed to pathogens, such as viruses and (myco)bacteria, and that such pathogenic exposure contributed significantly in the shaping of the modern human immune system (*15, 18, 19*). Human leukocyte antigen (HLA) class I and class II molecules are involved in antigen presentation to T-lymphocytes, which initiates the adaptive immune response. The heterogeneity of HLA genes (*HLA-A, -B, -C*) showed a strong correlation with SARS-CoV-2 infection susceptibility, severity and progression (*20*). It is estimated that the contribution of Neanderthal *HLA-A* alleles to non-Africans is 50-95% (*15, 21*). These findings point to a vital role of HLA molecules in host defense against viral infection and that these inherited HLA alleles provide an evolutionary advantage to modern human survival (*15*). A disadvantage of Neanderthal admixture may however be susceptibility to (auto)immune diseases associated with HLA. Archaic HLA alleles may have been positively selected and preserved in modern genomes because they were beneficial to the survival of Neanderthal and early modern humans encountering similar local pathogens at the time. However, some of the archaic alleles have been associated with development of autoimmune and auto-inflammatory diseases such as Behçet’s disease in modern humans (*22*). Furthermore, genetic variants inherited from Neanderthals have been shown to contain risk factors for a range of diseases including systemic lupus erythematosus, biliary cirrhosis, Crohn’s disease, and type 2 diabetes (T2D) (*23*). Recently, it was also shown that a gene cluster on chromosome 19, that introgressed from the Neanderthal lineage is a major risk factor for severe symptoms upon SARS-CoV-2 infection (*24*). These studies suggest that the gene flow from Neanderthals could also negatively affected modern humans.

Evidently, the gene flow between early modern humans and archaic humans has undergone strong balancing selection during the course of human evolution. So, we wondered whether genes that have strong detrimental consequences on the function of our immune system were present in Neanderthals and introgressed to the modern human genome. In modern humans the diseases associated with these errors in the immune system are called “inborn errors of immunity” (IEI). These include primary immunodeficiencies, autoimmune diseases and auto-inflammatory diseases. Currently, more than 450 IEI-associated genes that lead to a profound impairment of the immune system and therefore have a significantly impact on survival and quality of life are known (*25*). In this study we screened the genomes of three Neanderthals and one Denisovan for the presence of (potentially) damaging genetic variants in genes associated with IEI to explore our hypothesis that defects in the immune system might have contributed to the disappearance of the archaic humans.

## Methods

Variant call format files from the Neanderthal from the Denisova Cave in the Altai mountains (*Denisova 5*, referred to as Altai Neanderthal) were downloaded (http://cdna.eva.mpg.de/neandertal/altai/AltaiNeandertal/VCF/) per chromosome and uploaded in our local Galaxy cloud (*3, 26*). Files were filtered for variants (removing reference genotype 0/0) and merged. Variants (6,374,229) were annotated using ANNOVAR (*27*). The annotated variants file was exported and uploaded into TIBCO Spotfire Analyst 12.0.0 (https://www.tibco.com/) for filtering. The variants were filtered for IEI candidate genes (448 unique genes) based on the 2022 updated classification from the international union of immunological societies (IUIS) resulting in 76,516 variants (*25*). Subsequently, these variants were filtered for protein affecting variants (PAV): exonic and splicing variants (336 variants). Next, the variants that were absent or present in less than 0.01 in the control cohorts (GnomAD_exome, GnomAD_genome, 1K, ExAC, and GoNL) were selected (134 variants) and exported to excel. The variants were filtered on combined annotation dependent depletion (CADD v1.6) scores (*28*). Fifty-three variants had a CADD (v1.6) score ≥20 and 39 of these variants with a QUAL score (phred-scaled quality score) above 100 were selected. Among them, 24 variants were homozygous, and 15 variants were heterozygous in the Altai Neanderthal genome. All variants were checked in the genome browser from the Max Planck Institute (https://www.eva.mpg.de/genetics/genome-projects/ and https://bioinf.eva.mpg.de/-jbrowse/?loc=1%3A99708627..149558751&tracks=&highlight=) for the presence of the variants in the genomes of the Neanderthal from the Vindija Cave in Croatia (*Vindija 33.19* referred to as Vindija Neanderthal), and the Denisovan also from the Denisova Cave in the Altai mountains (*Denisova 3* referred to as Denisovan) (*3, 6, 29*). In addition, the presence of all variants were checked in the genome from the Neanderthal found in the Chagyrskaya Cave in the Altai Mountains by uploading the BAM files (http://ftp.eva.mpg.de/neandertal/Chagyrskaya/BAM/) (*30*) into our Galaxy cloud, and visualized by integrative genome viewer (IGV) version 2.3.94 (*31*). Five variants (*MTHFD1; WRAP53; NOS2; PEPD; CFTR*) in our homozygous or heterozygous lists were not found in the Max Planck Institute genome browser therefore these genes were removed from further analysis. Finally presence of the variants were checked in modern human genomes (Dai (HGDP01308), French (HGDP00533), Han (HGDP00775), Mandenka (HGDP01286), Mbuti (HGDP00982), Papuan (HGDP00546), San (HGDP01036), Sardinian (HGDP01076), Yoruba (HGPD00936), Karitiana (HGDP01015), Australian (WON,M), Australian (BUR,E), Mixe 0007 (MIXE 0007), and Dinka (DNK07)) from the Max Planck Institute genome browser and primate genomes (Chimp, Gorilla, Orangutan, Gibbon, Rhesus, Macaque, Baboon, Green Monkey, Marmoset, Squirrel Monkey, Bushbaby) from the UCSC genome browser. The human gene mutation database (HGMD) was used to check if the variants have been described before.

## Results

### Identification of variants in genes associated with IEI in Neanderthal and Denisovan genomes

To determine if Neanderthals and Denisovan genomes contained genetic variants in genes associated with IEI we used the variant call format files from a Neanderthal from the Altai mountains which contained 6,374,229 variants compared to the Homo sapiens genome assembly (hg19) (Figure 1A). From these variants, only the variants present in genes related with IEI were selected. These variants were filtered for protein affecting variants, the absence in control cohorts, with a CADD score equal or higher than 20, and with a QUAL score (phred-scaled quality score) above 100, leaving us with 39 variants in the Altai Neanderthal genome. Twenty-four of these variants were homozygous, and 15 variants were heterozygous (Figure 1A).

**Figure 1.**
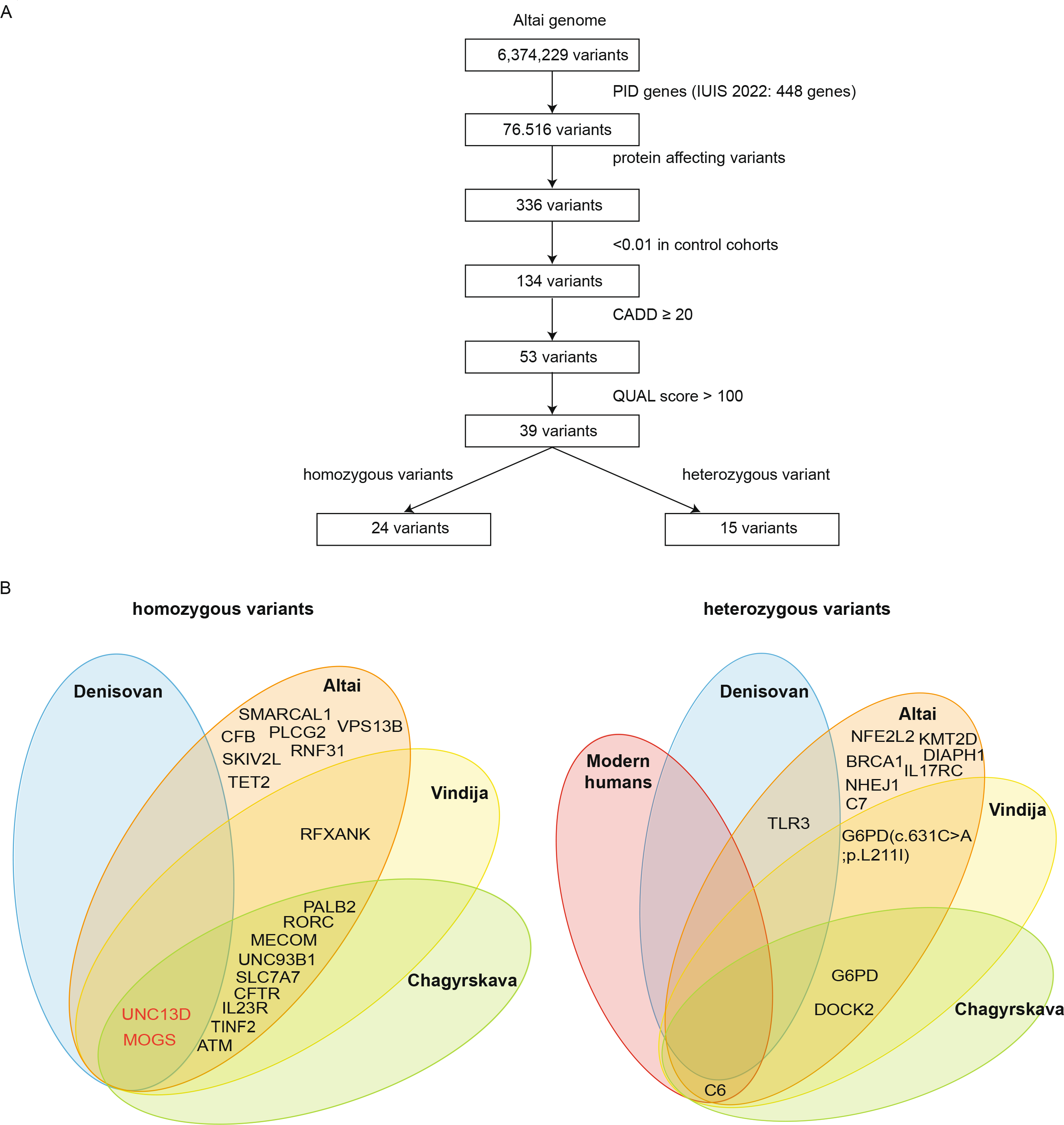
Genetic variants associated with IEIs. Filtering scheme for the identification of variants related to IEIs in archaic humans (Altai, Vindija, Chagyrskaya Neanderthals; Denisovan); modern humans and primate genomes (A). Venn diagrams of the 24 IEI-related homozygous (left panel) and 15 heterozygous (right panel) variants found in Altai, Vindija, Chagyrskaya Neanderthals; Denisovan; modern humans and primate genomes (B). Genes with variants that are common to all archaic humans (Altai, Vindija, Chagyrskaya Neanderthals; Denisovan) are highlighted in red.

Subsequently, we checked if these 24 homozygous and 15 heterozygous variants from the Altai Neanderthal were present in the published genomes of the Vindija Neanderthal, Chagyrskaya Neanderthal, and Denisovan genome. In addition, we checked if the variants were present in modern human genomes or primates. While none of the homozygous and heterozygous variants were found in the primate genomes, one variant (NM_001115131(C6):c.1848G>A,p.M616I) in the 15 heterozygous variants was also present in some modern genomes.

### Homozygous variants in IEI-related genes

Interestingly, we identified three variants in two different genes that were homozygous in all archaic humans genomes (Table 1). Both genes (*UNC13D* and *MOGS*) have been associated with an autosomal recessive form of IEI. Two of the variants were located in *UNC13D* gene NM_199242(UNC13D):c.1889C>T,p.A630V (rs372467471,CADD:26.2) and NM_199242(UNC13D):c.2828A>G,p.N943S (rs147748627, CADD:24.3). Both variants have very low allele frequency in the genome aggregation database (gnomAD) (0.000004063 and 0.005 respectively). Autosomal recessive variants in *UNC13D* cause hemophagocytic lymphohistiocytosis (HLH), a rare, life-threatening hyper-inflammatory immune disorder (*32*). HLH is characterized by uncontrolled activation of macrophages and T lymphocytes, and increased inflammatory cytokine production (cytokine storm) that leads to multi-organ failure. The *UNC13D* variants are in the Munc13-homology domain 1 (MHD1) and the Ca^2+^-binding (C2B) domain of Munc13-4 protein respectively (Figure 2A and Figure 3A). The *UNC13D* variant p.(Ala630Val) has not been previously reported in modern human genome, but the p.(Asn943Ser) variant has been described in 2014 in two patients presenting with HLH (*33*).

**Table 1.**
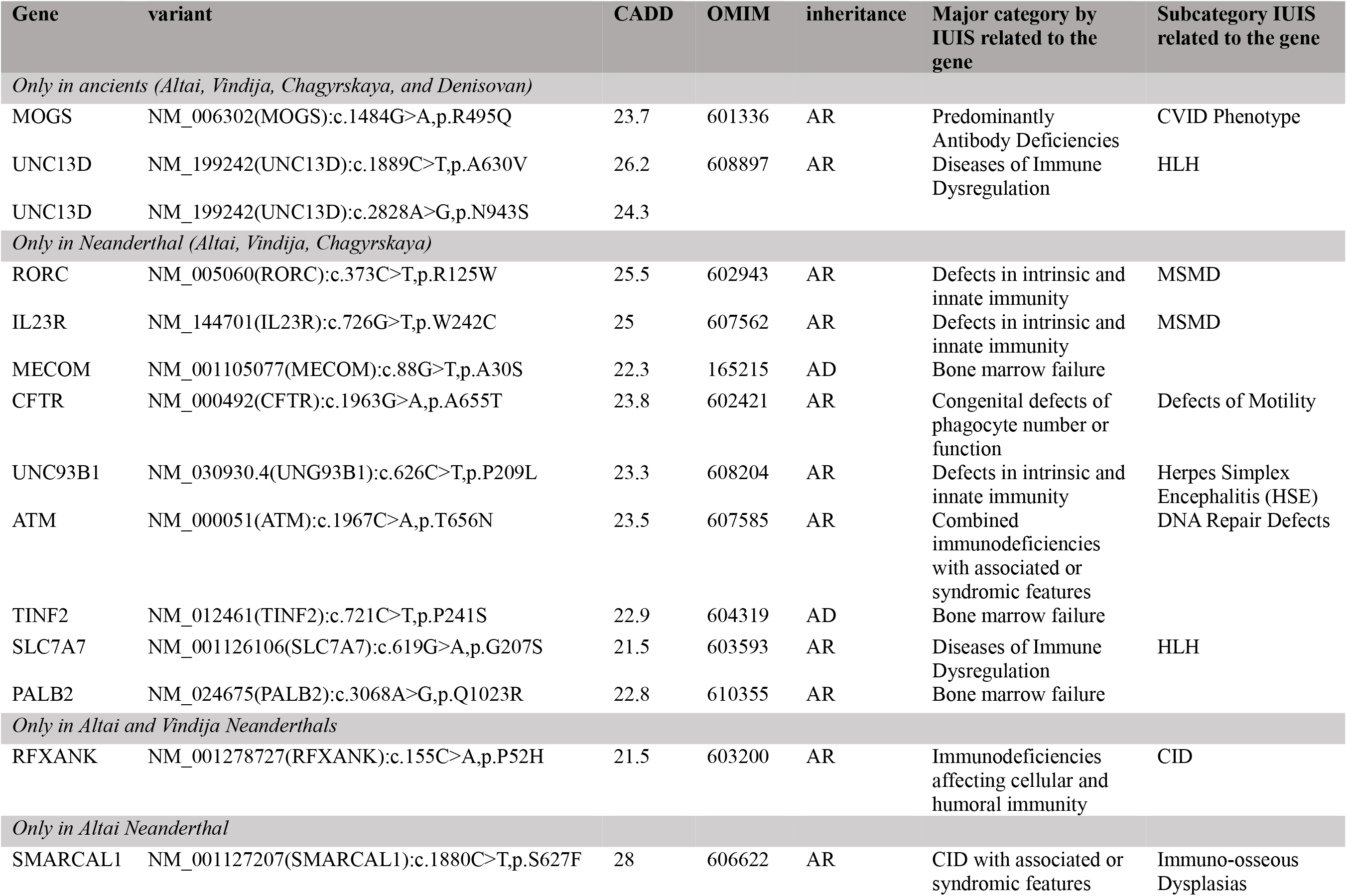
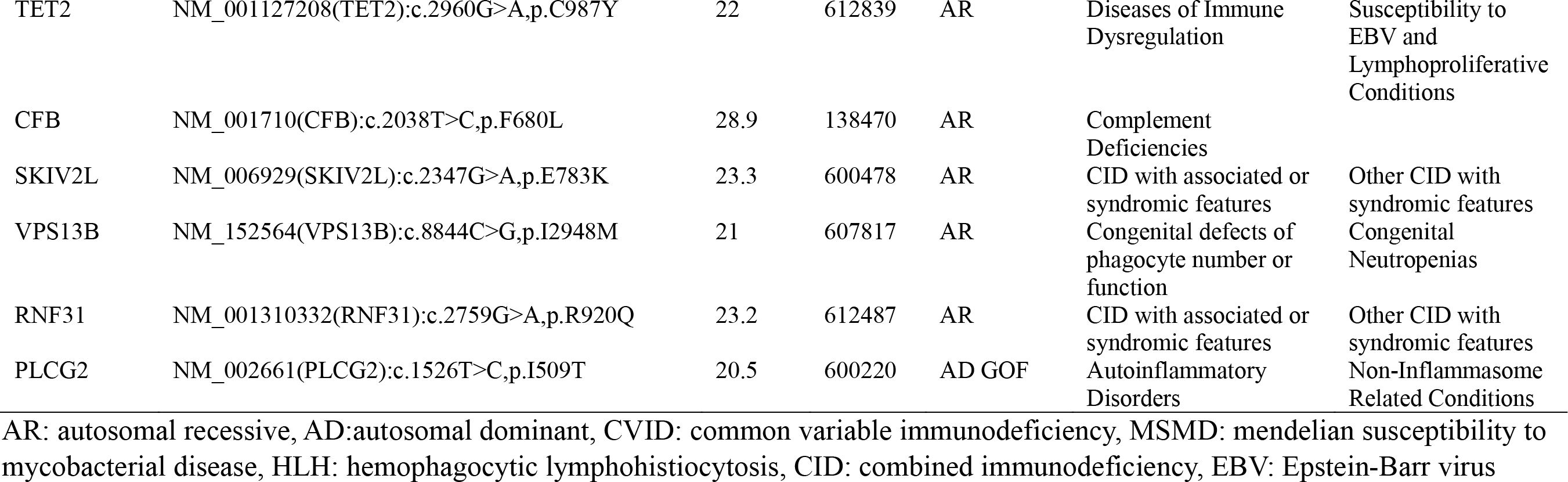
Homozygous variants in IEI-related genes in the archaic genomes

**Figure 2.**
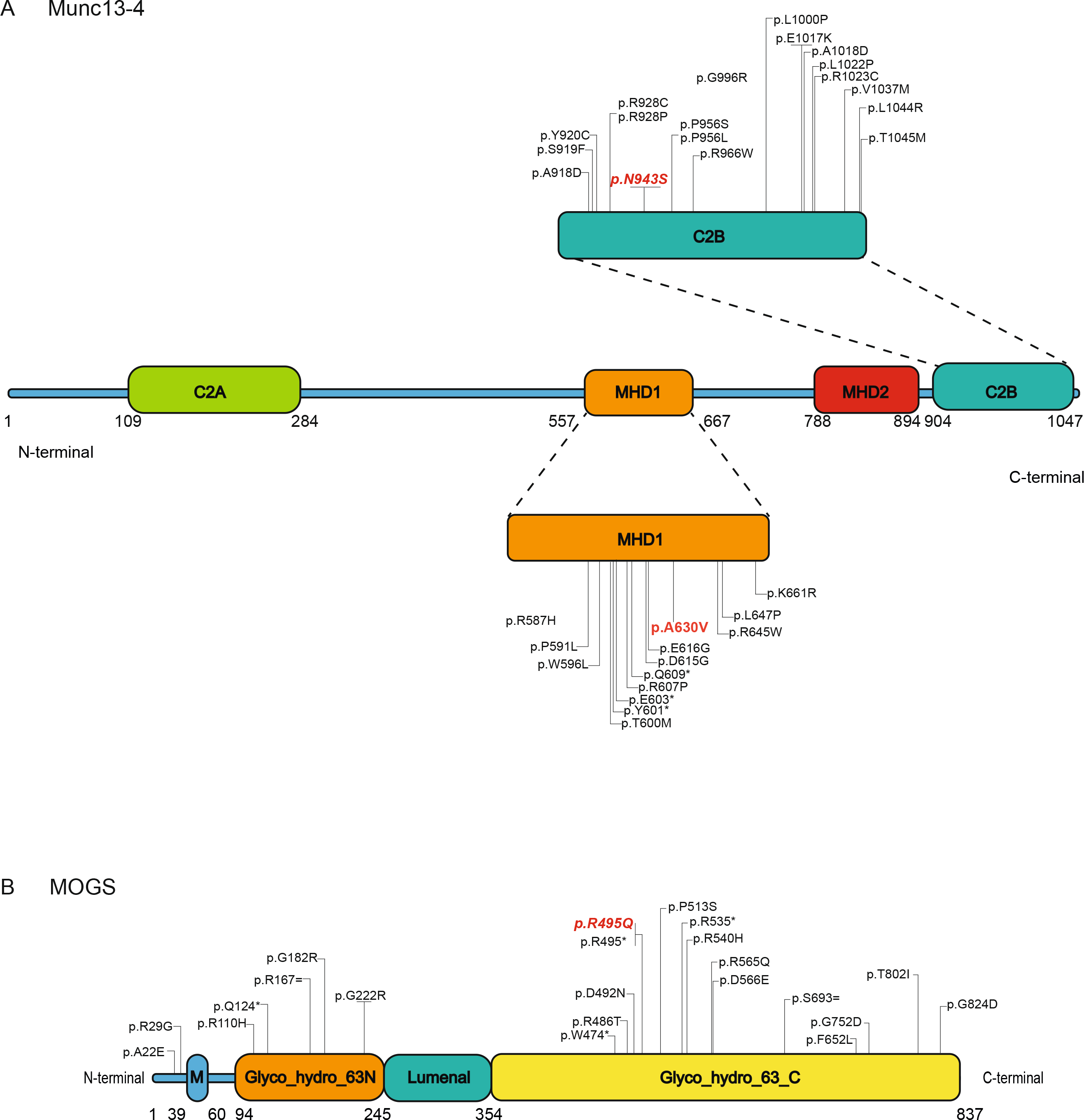
Schematic presentation of Munc13-4 and MOGS protein structure. Linear representation of the human Munc13-4 protein. Variants associate with HLH in the Munc13-homology domain 1 (MHD1) and the Ca^2+^-binding (C2B) domain are depicted (A). Linear representation of the human MOGS protein. Variants in the domain which provides Signal-anchor for type II membrane protein is marked M and in blue; the N-terminal beta sandwich domain found in glycosyl hydrolase family 63 proteins is marked Glyco_hydro_63N and in orange; the C-terminal catalytic domain found in glycosyl hydrolase family 63 proteins is marked Glyco_hydro_63C and in yellow (B). Variants described in modern humans are italicized; homozygous variants identified in archaic humans (Altai, Vindija, Chagyrskaya Neanderthals; Denisovan) is coloured red.

**Figure 3.**
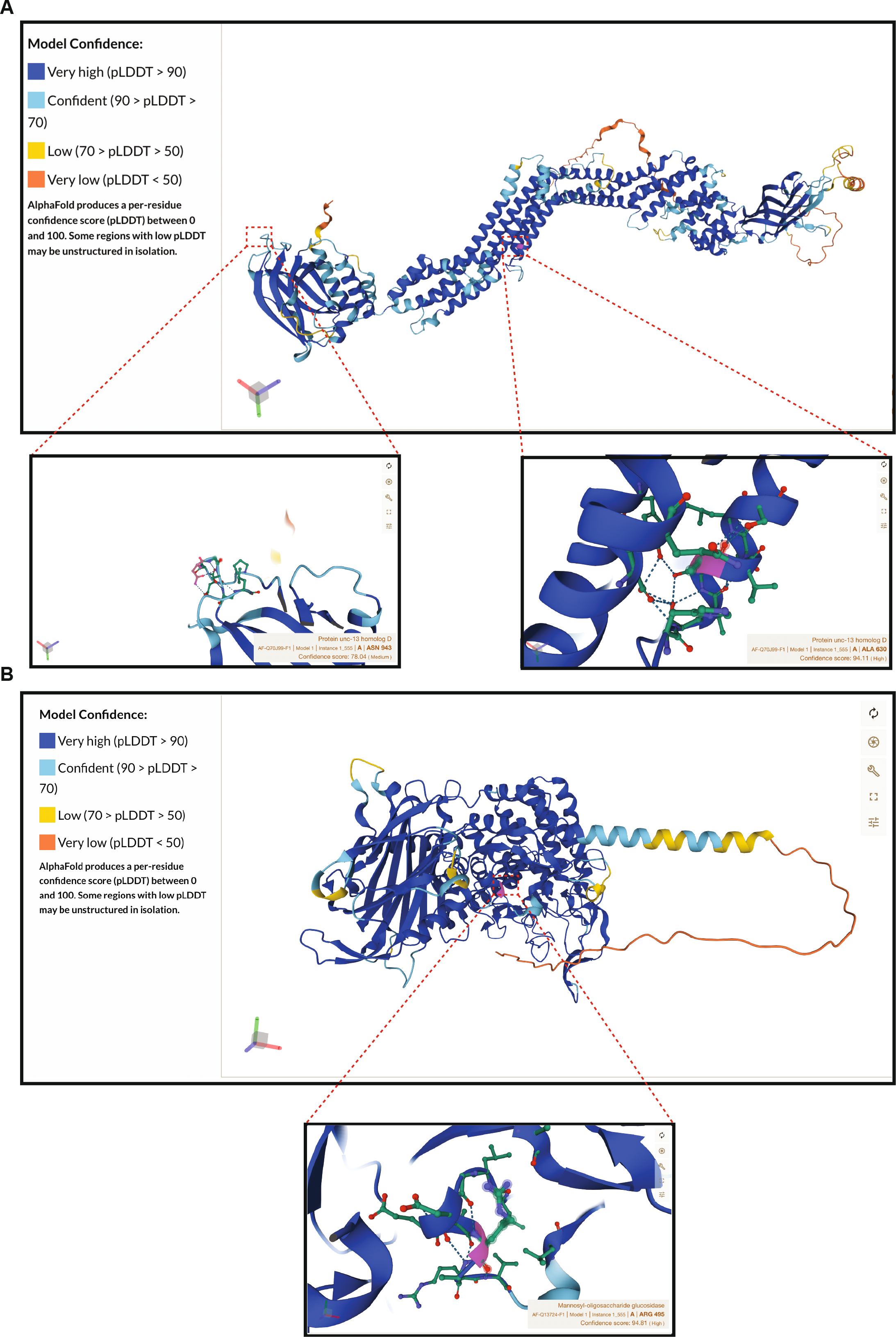
3D structure of human Munc13-4 and MOGS proteins. Predicted 3D structure of the human Munc13-4 protein extracted from structural data in UniProtKB. A magnified view of specific residue sites of identified variants of *UNC13D* (p.A630V and p.N943S) are highlighted below (A). Predicted 3D structure of the human MOGS protein extracted from structural data in UniProtKB. A magnified view of specific residue site of identified variant of *MOGS* (p.R495Q) is highlighted below (B).

The other variant that was present in all archaic genomes is located in the *MOGS* gene (Figure 2B and Figure 3B), which encodes for α-glucosidase 1, also known as mannosyl-oligosaccharide glucosidase (MOGS) (*34, 35*). Loss-of-function defects in *MOGS* underlie MOGS-CDG, a group of disorders caused by inborn errors of glycosylation, previously known as congenital disorders of glycosylation type II (CDG-IIb; OMIM;606056). Pathogenic variants of *MOGS* influence the processing of protein-bound glycans as well as glycoprotein folding and quality control (*36*). Additionally, *MOGS* is a known regulator of the HLA-class I antigen presentation pathway. Knocking-out of *MOGS* in murine cancer cell lines results in a significantly reduced expression of MHC-I (*37, 38*). The variant in the *MOGS* gene (NM_006302(MOGS):c.1484G>A,p.R495Q (rs34075781, CADD: 23.7)) is in the C-terminal catalytic domain (Figure 2B). This variant has a 0.0001 allele frequency in controls and was reported as a variant of unknown significance (VUS) in 2021 in a patient with common variable immunodeficiency (CVID) (*39*).

Nine homozygous variants were present in all Neanderthal genomes (Altai, Vindija, and Chagyrskaya), but not in the Denisovan genome. Most of these variants are of unknown significance because they have not been described before. However, the amino acid affected by the variant in the *ATM* gene (NM_000051(ATM):c.1967C>A,p.T656N) has been previously described to be affected in patients with Ataxia Telangiectasia (p.T656P) (*40*). One variant on a gene which is associated with HLA-class II deficiency (NM_001278727(RFXANK):c.155C>A:p.P52H) was shared by the Altai and Vindija Neanderthals but absent in the Chagyrskaya Neanderthal genome. Eight variants were only present in the Altai and Vindija Neanderthals, or even only in the Altai genome, indicating differential expression in archaic humans. None of these have been described before.

### Heterozygous variants in IEI-related genes

Fifteen variants in IEI-related genes were heterozygous in the Altai genome, but none of these were present in all archaic humans. However, three variants were present in all the Neanderthals (Altai, Vindija and Chagyrskaya). Interestingly, two of three variants are in the *G6PD* gene. The NM_001042351(G6PD):c.620T>C:p.M207TT variant has not been reported yet. The NM_001042351(G6PD):c.590A>T:p.Y197F variant in *G6PD* has also not been described, but another variant on the same amino acid (p.Y197S) has been described in boys with glucose-6-phosphate dehydrogenase deficiency (*41*). The third variant present in all Neanderthals is in the *DOCK2* gene (NM_004946(*DOCK2*):c.1855G>T:p.G619C). This gene has been recently reported to be involved in immune response to SARS-Cov2 infection and the development of severe COVID-19 (*42*). The inheritance of G6PD is X-linked, so the variants in *G6PD* could potentially affected the archaic males.

Most of the other heterozygous variants are related to autosomal recessive traits, so their clinical impact remains highly uncertain. However, the variant in Toll-like receptor 3 (*TLR3)* (NM_003265(TLR3):c.2276A>T:p.Y759F) shared by the Altai Neanderthal and Denisovan has an autosomal dominant inheritance. Loss-of-function disease-causing variants in *TLR3* have also been described in patients with herpes encephalitis (autosomal dominant and recessive) and severe COVD-19 infection due to impaired type I IFN induction (*43, 44*). The other two genes that are associated with autosomal dominant inheritance are the *KMT2D* and *NFE2L2* genes, which were only found in the Altai Neanderthal, and both are associated with combined immunodeficiency with syndromic features (Table 2). The exact variants in these genes have not been described, but a variant on the same amino acid as the p.A5212T variant in the *KMT2D* gene has been described in a patient with Kabuki syndrome (*45*).

**Table 2.**
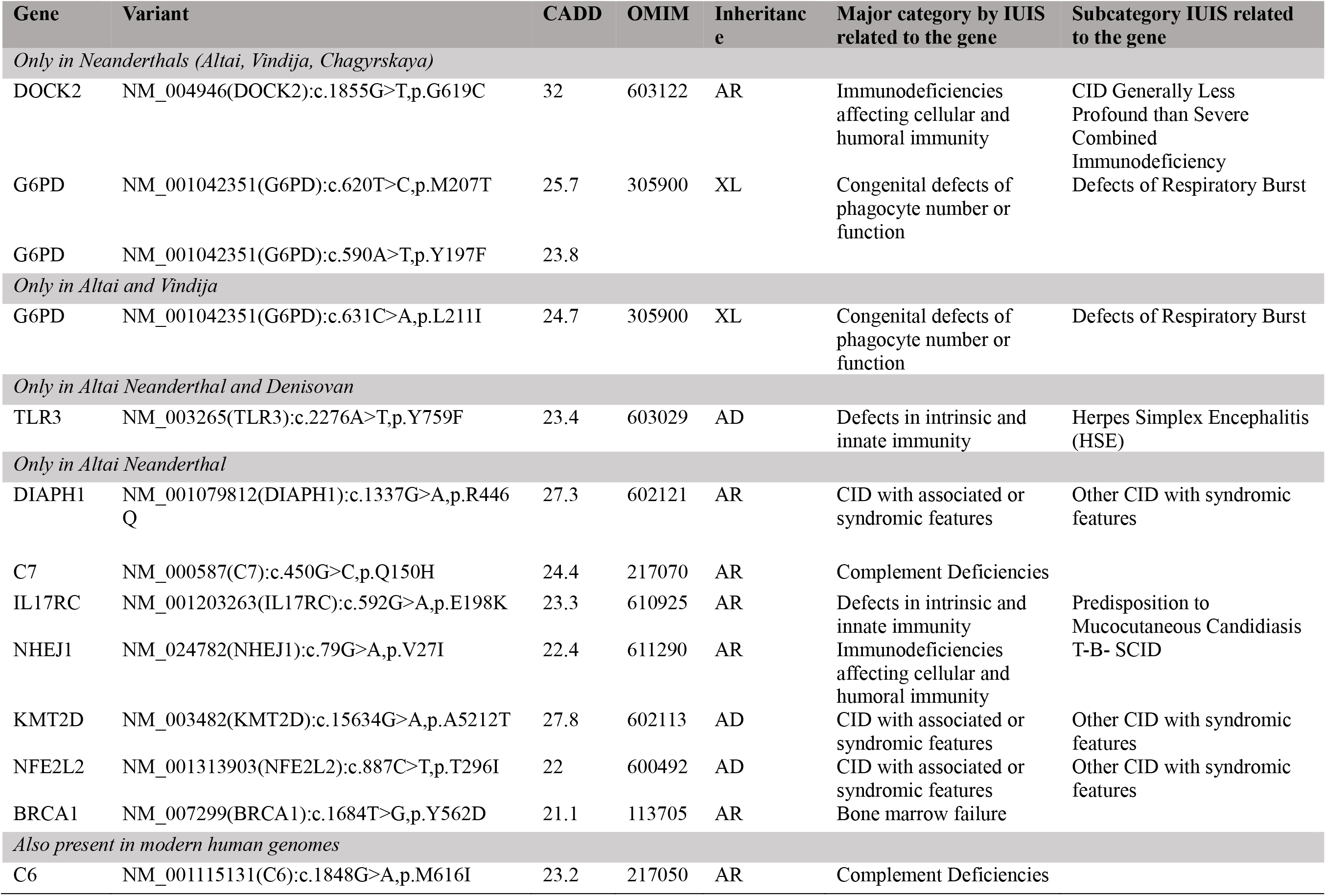
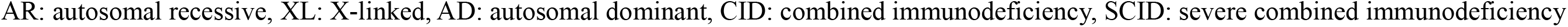
Heterozygous variant in IEI-related genes in the archaic genomes

## Discussion

In this study we investigated whether genetic variants that could have detrimental effects on the function of the immune system were present in the genomes of three Neanderthals and one Denisovan. We hypothesized that those variants, upon (specific) pathogenic exposure, could contribute to the extinction of the Neanderthal population. We first determined whether we could identify potentially damaging variants in the IEI-related genes in the genome of the Altai Neanderthal, and subsequently checked which of these variants were also present in the genomes of the Vindija Neanderthal, Chagyrskaya Neanderthal and the Denisovan. In total, we identified 24 homozygous and 15 heterozygous variants in the Altai Neanderthal from the human IEI gene panel (*25*). Most of these variants were not present in the other archaic human genomes studied, so it is unlikely that these variants were strong determinants in the extinction of Neanderthals. In contrast, we identified three homozygous variants in two different genes that were shared by all four archaic human genomes studied, while these variants were not present in the genomes of modern humans nor in the genomes of primates. This is interesting because the three Neanderthals and the Denisovan lived in different areas and time periods. The Altai Neanderthal and the Denisovan were found in the same Denisova Cave in the Altai Mountains in Siberia, but they were found in different sublayers. The Denisovan should be older than the Altai Neanderthal which was estimated to live 50,300±2,200 years ago (*3*). The Chagryrskaya Neanderthal was also found in a cave in the Altai mountains, but this cave was 100 km away from the Denisova cave and genetic analysis showed that she was more related to Neanderthals from western Eurasia (*30*). The archaeological layer in which the Chagurskaya Neanderthal was found, was estimated to be 60,000 years old. The Vindija Neanderthal was found in a cave in Croatia and is estimated to be 45,500 years old (*29*). That all three variants were present in all archaic genomes suggest that they must have been present before the different lineages diverged.

*UNC13D* encodes for Munc13-4, which is a crucial protein for the function of NK cells and cytotoxic T lymphocytes (CD8^+^) to destruct virus-infected cells and tumor cells, but also for the regulation of activated macrophages when a pathogen is cleared (*46*). Loss-of-function variants in *UNC13D* result in HLH, a complex inflammatory and often severe and fatal disease (*32, 46*). HLH is triggered by infections as NK-cell and cytotoxic T-cell dysfunction prevent efficient antigen clearance. Because of the defects in *UNC13D*, NK cells and cytotoxic T lymphocytes struggle with the successful elimination of pathogens, and in addition, fail to lyse the activated macrophages (*47*). This results in haemophagocytosis of host platelets, leucocytes and red blood cells. In addition, the (over)activation of the cytotoxic T lymphocytes, NK cells, and macrophages induces a cytokine storm in which large amounts of cytokines including IL-1β, IL-6, IL-18, IFN-γ, and TNF-α are produced and released (*48*). This cytokine storm is thought to cause multi-organ failure in HLH (*49*). Patients with HLH usually have poor prognosis (*50, 51*), with the current treatments, mortality rates among HLH in adults are around 60% (*52*).

The two homozygous *UNC13D* variants that were only found in the archaic human genomes were in the MHD1 region (c.1889C>T:p.A630V) and in the C2B region (c.2828A>G:p.N943S) in the Munc13-4 protein. Both variants are reported in control cohorts, however at a very low allele frequency (c.2828A>G:p.N943S; allele frequency: 0.0006630, c.1889C>T:p.A630V; allele frequency: 0.000007362). The c.1889C>T variant has not been described in patients with HLH before. Nonetheless, the c.2828A>G variant was described in two patients with HLH (*33*). These patients were heterozygous for this variant, and expressed an additional heterozygous variant in another gene associated with HLH, suggesting a digenic mode of inheritance that results in a synergistic defect (*33*). It is noteworthy that the two variants in *UNC13D* did not introgress into the modern humans, implying that these variants may have a negative effect on the immune system of archaic humans after exposure to certain microbes. Although there are not patients with HLH described who are homozygous for one of the two *UNC13D* variants identified in our study, the data so far suggests that it is very likely that both variants, and especially the c.2828A>G variant could have impaired the function of Munc13-4 in the archaic humans.

HLH in patients with a defect in *UNC13D* is usually triggered by an infection. Infections with *mycobacterium tuberculosis* (*Mtb*), Epstein-Barr virus, Herpes simplex virus and Varicella zoster virus have been described to trigger HLH (*53–57*). In Neanderthal DNA, fragments of various infectious agents were detected in dental pulp from Neanderthals, including bacteria from the Actinobacter phylum (*58*). One of the most studied bacteria from the Actinobacteria phylum is *Mtb*. *Mtb* is one of the most successfully bacterial species that spread globally. Currently, more than 1 billion modern humans are infected. Individuals infected with *Mtb* have a 5–10% lifetime risk of getting clinically active tuberculosis. One person with tuberculosis can yearly infect 5–15 other people through close contact. Without treatment, almost half of the patients with HIV-negative tuberculosis on average will die (*59*).

The evolutionary timing and spread of the *Mtb* was reported by Wirth et al, using mycobacterial tandem repeat sequences as genetic markers (*60*). Using Bayesian statistics and experimental data on the variability of the mycobacterial markers in infected patients, they estimated the age of the *Mtb* at 40,000 years. This coincides with the expansion of early modern human populations out of Africa. An interesting hypothesis could be that migration of early modern humans out of Africa carried along *Mtb*, spreading the pathogen across Eurasia, thus exposing archaic humans such as Neanderthals to this “new” pathogen. As indicated previously, induction of HLH needs a trigger. This was also shown in a mouse model of HLH. The mice that had a donor splice site in *Unc13d*, the mouse orthologue of human MUNC13-4, did not spontaneously developed an HLH phenotype, but only upon a specific infectious trigger (*61*). It could thus be that “new” pathogens like *Mtb* were the specific trigger that caused an HLH phenotype in the archaic humans.

Glycosylation is one of the most fundamental protein modification processes in all living organisms as the majority of secreted and membrane-bound proteins are glycoproteins. An estimated proportion of 2% of human genes encode for glycosylation-related proteins (*62*). N-glycosylation refers to the attachment of glycans to the nitrogen atom of an asparagine residue in a protein, whereas O-glycosylation is the attachment of glycans to the oxygen atom of serine or threonine residues (*63*). N-glycosylation is the most common co-translational/post-translational protein modification in human cells (*64*). MOGS is expressed ubiquitously in the endoplasmic reticulum, where it functions as the first enzyme involved in the trimming of N-glycans in the N-glycosylation pathway. This initial step is crucial to the further processing of N-glycans. When the enzyme is disrupted various important biological processes are impaired, including protein folding, stability and transition. Genetic loss-of-function variants in the *MOGS* gene causes the extremely rare congenital disorder of glycosylation type IIb (CDG-IIb), also known as MOGS-CDG. MOGS-CDG presents as a paradoxical immunologic phenotype associated with severe hypogammaglobulinemia, in absence of overt clinical indications of infection along with various neurologic complications (*65*). The mechanism behind the hypogammaglobinemia was an unexplained shorter immunoglobulin half-life. A resistance to infections such as HIV and influenza viruses could be explained by impaired viral replication and cellular entry of these enveloped viruses which are heavily glycosylated (*65*). Glycoproteins are also important for microbes itself. The SARS-CoV2 spike protein, for instance, is an extensively glycosylated protein which protrudes from the viral surface and mediates host-cell entry by binding to angiotensin-converting enzyme 2 (ACE2) (*66*). On the other hand, glycosylation is also a factor in the immune defense against microorganisms and pathogens. Numerous effector systems of the immune system, including cytokines, cytokine receptors (*67*), and HLA require fine-tuned glycosylation in order to function optimally (*68*).

In 2014, Pääbo et al. reported the same *MOGS* variant ((NM_006302):c.1484G>A,p.R495Q (rs34075781; CADD: 23.7) in relation to bone morphology, abnormalities in palates in the genomes of three Neanderthals (El Sidrón, Vindija, and Altai genomes) (*7*) The variant is located in the C-terminal catalytic domain which catalyzes the cleavage of N-glycans. This variant is ultra-rare (with allele frequency of up to 0.001%) in control cohorts, and it was previously described once in a CVID patient, as a heterozygous VUS. MOGS is one the key genes regulating HLA-I expression and plays a significant role in immune defense by regulating immunoglobulin half-life, viral replication, and cell entry. The absence of the homozygous p.R495Q variant in the modern human population, suggests that this variant may be damaging upon exposure to specific pathogens.

The relationship between archaic alleles and disease in modern humans remains unclear. Since 2019, the SARS-Cov2 pandemic has provided evidence linking the disease severity to genes inherited from Neanderthals (*24, 69*). In South Asians, especially Bangladeshis, a Neanderthal haplotype of a gene cluster on Chromosome 3 is adaptively retained and is linked to a more severe response to Sars-CoV-2 infection (*24*). Increased levels of a Neanderthal haplotype of 2′5′ oligoadenylate synthase (OAS1) in Europeans, on the other hand, confers protection against Sars-CoV-2 infection (*69*).

In conclusion, our study is the first to specifically describe the presence of IEI-related gene-variants in the genomes of Neanderthals and Denisovian. We discovered three variants on two genes that are linked to rare IEI disorders in modern humans. All three of these variants were present in all the explored genomes of Neanderthals and Denisovan, but absent from modern human genomes (or were found heterozygous and in extremely low allele frequencies) and other primate genomes. This observation strongly suggests a deleterious effect of these variants on the immune system of archaic humans. Neanderthals and Denisovans survived a long period of time until, shortly after (by evolutionary standards) the Paleolithic migrations of modern humans (Homo sapiens). Studies by Houldcroft et al, indicated that infectious diseases are tens of thousands of years older than previously believed and thus co-evolved with humans for long time (*70*). They hypothesized that spreading of pathogens out of Africa could have been catastrophic to Neanderthal populations in Eurasia. Our findings add a novel layer to this with the identification of archaic-human-specific variants in genes associated with IEI in modern human. We therefore postulate that: 1). these archaic-human-specific IEI gene-variants were very detrimental and caused severe and fatal disease upon encounter of pathogens brought along by modern human, 2). Such unfortunate host (gene)-pathogen interactions may have expedited the extinction of Neanderthals.

## Acknowledgements

The research for this manuscript was performed within the framework of the Erasmus Postgraduate School Molecular Medicine. This work was supported by the China Scholarship Council for funding PhD fellowships. (No.201908440363 Z.Z)).

## Disclosure of conflicts of interest

The authors have no conflict of interest.

## Notes

http://cdna.eva.mpg.de/neandertal/altai/AltaiNeandertal/VCF/

https://www.eva.mpg.de/genetics/genome-projects/

https://bioinf.eva.mpg.de/jbrowse/?loc=1%3A99708627..149558751&tracks=&highlight=

http://ftp.eva.mpg.de/neandertal/Chagyrskaya/BAM/

### Competing Interest Statement

The authors have declared no competing interest.

